# Flexibility of systemic one-carbon metabolism partially buffers dietary methyl donor deficiency

**DOI:** 10.64898/2026.02.05.703880

**Authors:** Edmund D. Kapelczak, Rashel Jacobo, Sofia de Lourdes Mirabal, Katherine A. Dang, Vincent Hernandez, Tara TeSlaa

## Abstract

Choline is a methyl-rich nutrient used in lipid synthesis and catabolized to support one-carbon metabolism. Choline consumption in most humans remains less than the suggested adequate intake, yet how systemic metabolism compensates for choline deficiency is not fully described. Here, we use *in vivo* stable isotope tracing to explore the fate of choline in mammalian tissues. We find that choline is catabolized in the liver to support both the methionine cycle and the mitochondrial folate cycle in addition to its role in lipid synthesis. When dietary methyl donors are deficient, surprisingly, we find maintained systemic choline and methionine fluxes, but diminished contribution of choline to the folate cycle. To compensate for dietary methyl deficiency, systemic flux of serine is doubled by increased kidney synthesis which supplies one-carbon units for increased methionine synthesis in the liver. Our study suggests that systemic one-carbon flexibility can compensate for nutritional methyl deficiency by inter-organ nutrient exchange.

## Introduction

Choline is a semi-essential nutrient in humans that serves as a lipid head group and contains three methyl groups that can be transferred into one-carbon metabolism to support various biosynthetic pathways. While the body can produce choline through a series of three methylation reactions converting phosphatidylethanolamine (PE) into phosphatidylcholine (PC) by the enzyme phosphatidylethanolamine methyltransferase (PEMT), a complete deficiency of exogenous choline results in liver damage^1,2^. Despite its clear importance, most people consume less than the daily recommended amount of choline, and an individual’s personalized choline need can vary with genetic background, pregnancy, lactation, and disease^3–5^. Prolonged insufficient intake of choline may contribute to metabolic dysfunction-associated steatotic liver disease (MASLD) and cognitive decline in adults^3^.

Dietary choline comes in the form of either free choline or acyl-bound phosphatidylcholine (PC), which are often interconverted in the body. Catabolism of free choline into betaine can feed one methyl group into the methionine cycle, generating dimethylglycine, which can be used sequentially to donate the other two methyl groups into the mitochondrial folate cycle. The liver and kidney have the highest expression of enzymes in the choline catabolism pathway^6^, but how much choline contributes to one-carbon metabolism in these and other tissues has not been fully characterized. Recent studies using oral betaine have shown promise in treating patients with MASLD^7^, highlighting the potential importance of the catabolic fate of choline rather than its role in lipid synthesis.

Humans given total parenteral nutrition require choline supplementation to prevent liver damage^1^. Consistent with this, deficiency of dietary choline in rodents causes the accumulation of fat and progressive liver damage over the course of weeks to months^8^. In mice, dietary methionine is often removed in combination with choline to model MASLD since methionine donates methyl groups for PC synthesis by PEMT. When fed the methionine and choline-deficient (MCD) diet, decreased export of triglycerides from the liver in very low density lipoproteins (VLDL) particles is thought to be caused by decreased PC/PE ratio, leading to steatosis, which progresses to inflammation and fibrosis^7,9–11^. Despite liver damage, mice can survive for long periods of time on the MCD diet, suggesting that the body can compensate for severe methionine and choline deficiency.

Here, we investigate the contribution of choline catabolism to one-carbon metabolism and lipid synthesis in normal physiology. To do this, we use *in vivo* stable isotope tracing in conscious mice and investigate the fate of choline-, methionine-, and serine-derived one-carbon units. To gain a deeper understanding of the importance of choline, we then measure how these pathways respond to choline deficiency induced by the MCD diet.

## Results

### Choline feeds into the methionine cycle in liver and white adipose tissue

The methionine cycle produces S-adenosylmethionine (SAM), which is the universal methyl donor for most of the methylation reactions in the cell. Transfer of the SAM methyl group produces S-adenosylhomocysteine (SAH), which can then be converted to homocysteine, which can either be recycled for re-methylation to produce a new methionine or can be used in the transsulfuration pathway. Circulating methionine itself can feed into the methionine cycle, but if demand for methyl groups exceeds methionine availability, alternative methyl donors are thought to contribute. One alternative methyl donor is choline, which can be converted into betaine and donate a methyl group to homocysteine via the betaine homocysteine methyltransferase (BMHT) reaction (Figure 1a). Thus, we sought to quantify the contribution of circulating methionine and circulating choline to the methionine cycle in tissues. To assess circulating methionine’s contribution to the methionine cycle, we performed *in vivo* stable isotope tracing with [methyl-^13^C_1_]methionine and measured labeling of SAM (Fig. 1b).

**Figure 1.**
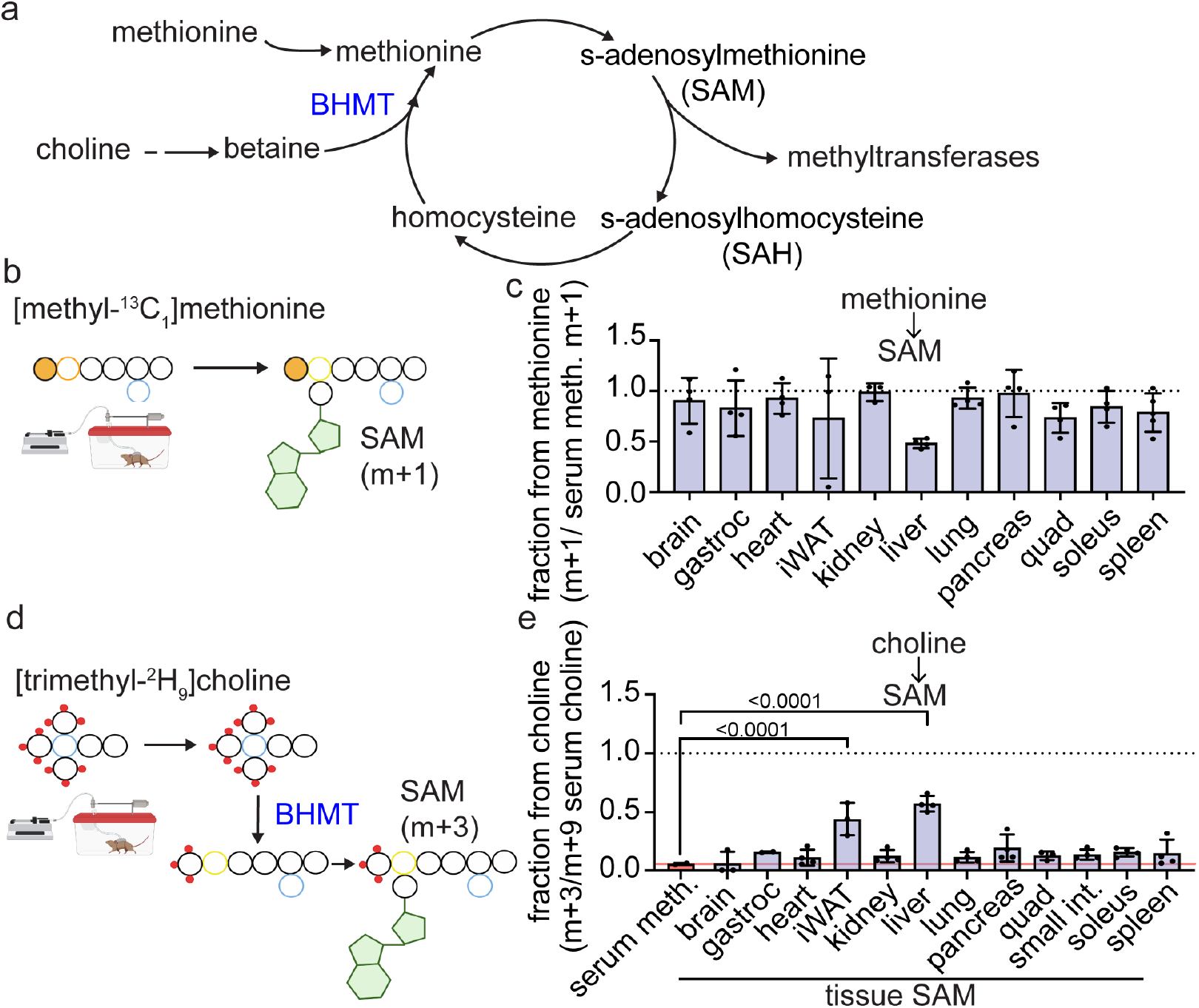
Methionine supplies methyl groups for all tissues, while choline is a unique substrate for liver and white adipose. (a) Schematic of the methionine cycle and nutrient methyl donor inputs. (b) Intravenous infusion of [methyl-^13^C_1_]methionine to measure the contribution of circulating methionine to tissue s-adenosylmethionine (SAM) pools. (c) Labeling of tissue SAM normalized to serum enrichment of methionine after 8 hour intravenous infusion of [methyl-^13^C_1_]methionine representing the fraction of SAM derived from circulating methionine. The dotted line indicates a labeled fraction of 1, which would imply complete labeling of the tissue SAM pool from methionine. (d) Intravenous infusion of [trimethyl-^2^H_9_]choline to measure the contribution of circulating choline to tissue s-adenosylmethionine (SAM) pools. (e) Labeling of tissue SAM normalized to serum enrichment of choline after 8 hour intravenous infusion of [trimethyl-^2^H_9_]choline representing the fraction of SAM derived from circulating choline. Dotted line indicates a labeled fraction of 1, which would imply complete labeling of the tissue SAM pool from choline. The red line represents the fraction of serum methionine that is labeled. Labeling below this line indicates serum methionine as a potential source of labeling rather than tissue autonomous production. Bars represent mean ± S.D. Each data point represents data from an independent mouse. P-values were calculated with ordinary one-way ANOVA with Dunnett correction for multiple comparisons. All experiments were performed in male *ad lib* fed C57Bl/6N mice during the light cycle. BHMT, betaine homocysteine methyltransferase

Isotopically labeled methionine was administered via continuous intravenous infusion at a constant rate for a duration of 8 hours in awake mice during the light cycle during which serum enrichment of the labeled methionine was measured (Extended Data Fig.1a). SAM pools in tissues were almost completely labeled indicating that most tissues use circulating methionine rather than relying on alternative methyl donors.

Notably, in the liver less than half of the SAM was labeled from circulating methionine, suggesting the presence of an alternative nutrient source contributing to the remaining SAM synthesis (Fig. 1c). Consistent with this observation, labeling of 5-methylthioadenosine (MTA), a SAM-derived metabolite in methionine salvage, was high in all tissues except the liver (Extended Data Fig. 1b). We could also observe labeling on products of methyltransferase enzymes such as choline from PEMT in the liver and serum and creatine from guanidinoacetate methyltransferase (GAMT) in the pancreas and serum (Extended Data Fig 1c-e). These data suggest that most tissues rely on exogenously supplied methionine for SAM production and that the liver is unique in that it makes SAM from other sources.

Betaine, which can be derived from choline, is used by betaine homocysteine methyltransferase (BHMT) to transfer a methyl group from betaine to homocysteine to form methionine. To investigate the contribution of choline to methionine and S-adenosylmethionine (SAM) synthesis, we next sought to infuse [trimethyl-^2^H_9_]choline. Because there were no published studies that we could find in literature where stable isotope-labeled choline was infused in mice, we first measured the choline rate of appearance to determine an infusion rate. We found that an infusion rate of 1.5 nmol · min^-1^ · g bodyweight^-1^ produced about 22% serum enrichment (Extended Data Fig. 2a), which corresponds to a rate of appearance of choline of about 5.6 nmol · min^-1^ · g bodyweight^-1^ (Extended Data Fig. 2b), which is in the range of many essential amino acids, suggesting moderately high turnover of circulating choline^12–14^.

**Figure 2.**
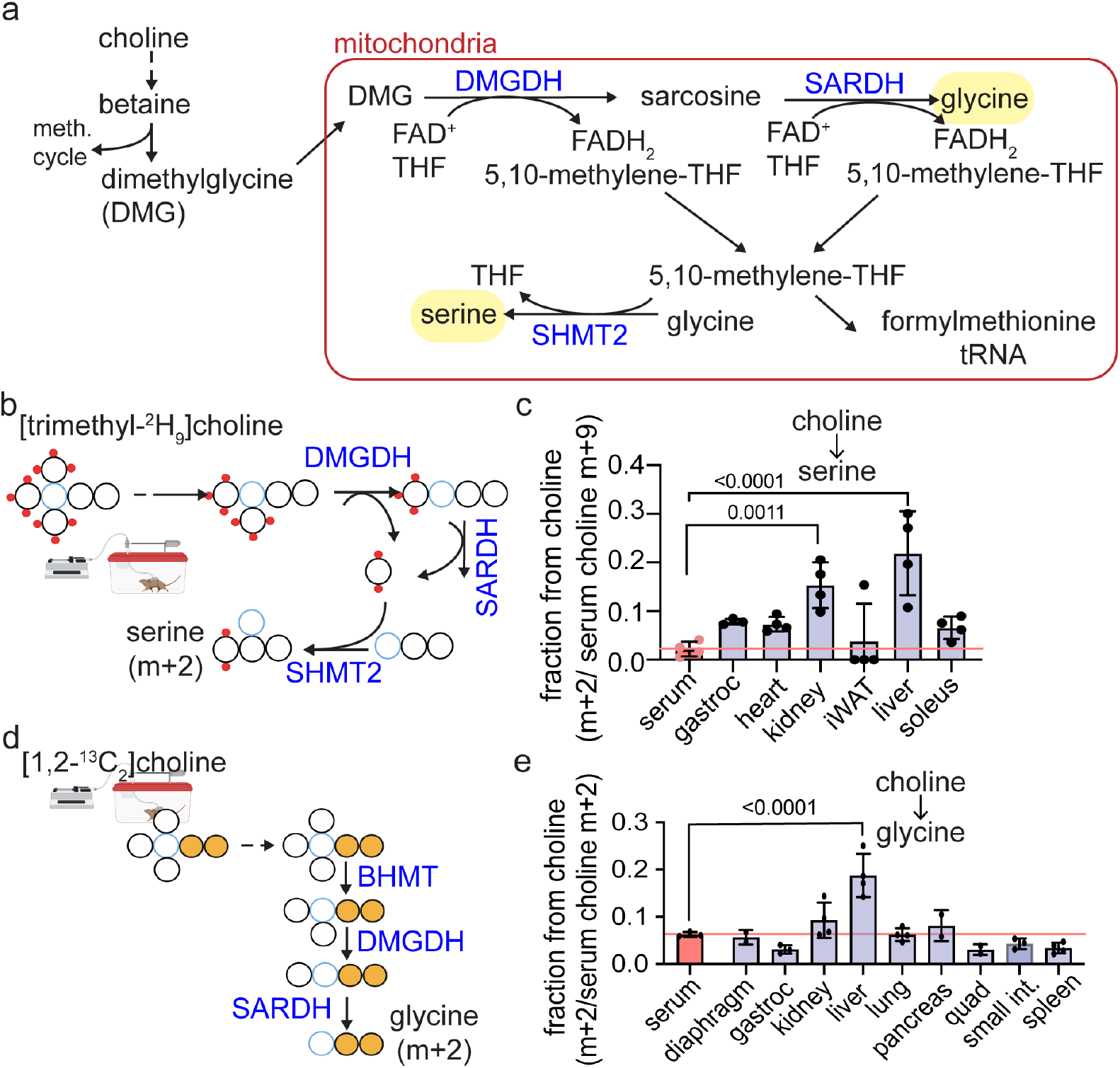
Choline supports mitochondrial folate cycle in liver. (a) Schematic showing the steps of choline catabolism that contribute to the mitochondrial folate cycle. (b) Intravenous infusion of [trimethyl-^2^H_9_]choline to measure the contribution of circulating choline to tissue serine pools. Because SHMT2 operates in reverse in the liver and kidney, serine serves as a readout of the mitochondrial folate cycle. (c) Normalized labeling of tissue serine from 8 hour intravenous infusion of [trimethyl-^2^H_9_]choline. The red line represents the fraction of serum serine that is labeled. Labeling at or below this line indicates that labeling is potentially sourced from serum serine pools rather than tissue autonomous production. (d) Intravenous infusion of [1,2-^13^C_2_]choline to measure the contribution of circulating choline to tissue glycine pools. (e) Normalized labeling of tissue glycine from 8 hour intravenous infusion of [1,2-^13^C_2_]choline. The red line represents the fraction of serum glycine that is labeled. Labeling at or below this line indicates that labeling is potentially sourced from serum glycine pools rather than tissue autonomous production Bars represent mean ± S.D. Each data point represents data from an independent mouse. P-values were calculated with ordinary one-way ANOVA with Dunnett correction for multiple comparisons. All experiments were performed in male *ad lib* fed C57Bl/6N mice during the light cycle. DMGDH, dimethylglycine dehydrogenase; SARSDH, sarcosine dehydrogenase; SHMT2, serine hydroxymethyltransferase

We next measured the contribution of choline to the methionine cycle by infusing [trimethyl-^2^H_9_]choline for 8 hours, which resulted in equal labeling of choline and betaine, suggesting that in mice on normal chow, there are no alternative sources of betaine other than circulating choline (Fig. 1d and Extended Data Fig 2c). Serum had only minimal labeling of circulating methionine, and most tissues had minimal labeling of SAM (Fig. 1d and Extended Data Fig. 2c-d). However, in the liver and white adipose tissue, we observed significantly higher labeling of SAM than circulating methionine, suggesting use of BHMT to make methionine within these tissues rather than uptake of labeled methionine from circulation (Figure 1e). Use of BHMT in the liver was supported by labeling of MTA and creatine (Extended Data Fig 2e-f). In contrast, we were not able to confirm the use of BHMT in adipose because labeling was undetectable, potentially due to low levels of MTA in the tissue.

Because choline only contributed to the methionine cycle in the liver and maybe the adipose, we also measured choline uptake and phosphorylation, which is the first step in the Kennedy pathway. Uptake and phosphorylation of choline was observed broadly across tissues, as indicated by labeling of phosphocholine (Extended Data Fig. 2g-h). Thus, choline is transported and phosphorylated in many tissues, but choline-derived betaine is only catabolized in the methionine cycle in the liver and white adipose. Overall, choline and methionine account for the majority of SAM production in mouse tissues, suggesting limited contribution from other sources, and choline is a unique source of SAM in the liver.

### Choline supplies one-carbon units for the liver mitochondrial folate cycle

Further downstream in the choline catabolic pathway after BHMT, dimethylglycine is used by dimethylglycine dehydrogenase (DMGDH) in the mitochondria to donate and oxidize a methyl group, producing sarcosine and 5,10-methylene-THF from THF and reducing an FAD^+^ to FADH_2_. Sarcosine is then used by sarcosine dehydrogenase (SARDH) for another oxidative demethylation to produce glycine, producing another 5,10-methylene-THF. This series of demethylation reactions produces two one-carbon units in the mitochondria in the form of 5,10-methylene-THF which can be used for local production of N-formylmethionine tRNA or nucleotides (Fig. 2a)^15,16^. These one-carbon units are critical for mitochondrial biosynthetic functions; in particular, formylmethionine-tRNA production is required for mitochondrial translation^16,17^.

To investigate the capacity of choline to contribute to the mitochondrial folate cycle, we sought to find readouts of the folate cycle using the [trimethyl-^2^H_9_]choline tracer. Our tracing strategy, which can specifically look at mitochondrial one-carbon units, is dependent on the directionality of the mitochondrial isoform of serine hydroxymethyltransferase, serine hydroxymethyltransferase 2 (SHMT2). In many cells, SHMT2 produces one-carbon units by catabolizing serine to glycine and 5,10 methylene-THF^15,18^, but in the liver and kidney, SHMT2 runs in reverse using one-carbon units from 5,10-methylene-THF to produce serine from glycine^19,20^ (Fig. 2a-b). Dimethylglycine and sarcosine produced from [trimethyl-^2^H_9_]choline have deuteriums on all methyl groups and, if used in the DMGDH and SARDH reactions, respectively, produce a M+2 5,10-methylene-THF with one deuterium transferred to form FADH_2_ (Fig. 2a-b). Thus, if choline feeds into the mitochondrial folate cycle in the liver and kidney, M+2 5,10-methylene-THF will be used to make serine by SHMT2 and we will observe M+2 serine (Fig. 2b). Labeling from [trimethyl-^2^H_9_]choline to serine in tissues was highest in liver with both kidney and liver showing significantly more labeling when compared to serum, suggesting that choline catabolism significantly contributes to the mitochondrial one-carbon pool in these tissues (Fig. 2c).

For a more comprehensive analysis of the DMGDH and SARDH reactions across tissues that do not exhibit reverse SHMT2 flux, we infused [1,2-^13^C_2_]choline, which has labeling on the choline backbone rather than the methyl groups (Fig 2d). Infusion of this choline tracer also resulted in equal labeling of the betaine pool when compared to choline, and equal labeling was also observed in circulating dimethylglycine, suggesting complete derivation of these nutrient pools from choline (Extended data Fig 3). Circulating glycine, on the other hand, was minimally labeled after 8 hours of infusion. Next, we analyzed tissue data for labeled glycine, which would support activity of DMGDH and SARDH to produce glycine from choline (Fig. 2d). We observed significantly more labeling of glycine in the liver when compared to serum (Fig. 2e). Thus, we conclude that the labeling observed in glycine in the liver is due to internal choline catabolism and not from taking up labeled glycine from the blood. These studies are consistent with previous studies that showed that liver mitochondria can respire when provided choline or dimethylglycine at rates similar to when they are provided with pyruvate and malate^21^. Combined, these data suggest that there is a unique role for complete choline catabolism in the liver, which is in contrast to the broader importance of choline’s contribution to the Kennedy pathway across all tissues.

**Figure 3.**
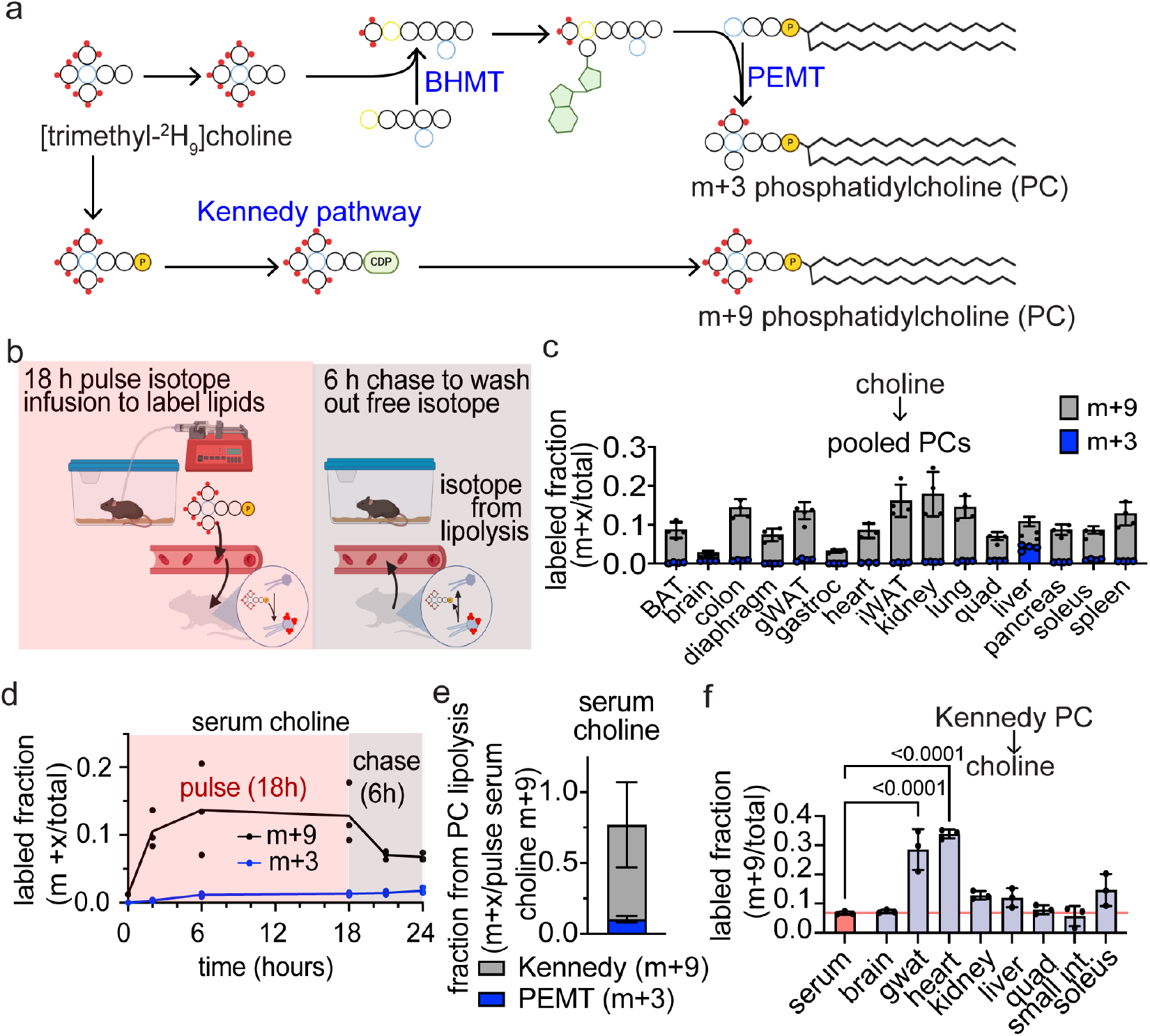
Free choline is recycled from lipids. (a) Tracing of [trimethyl-^2^H_9_]choline into phosphatidylcholine (PC) species. (b) Schematic of pulse chase infusion to measure the contribution of PC lipolysis to free choline pools. (c) Labeled fraction of pooled PCs at the end of the pulse chase infusion. (d) Labeled fraction of serum choline throughout the pulse chase intravenous infusion of [trimethyl-^2^H_9_]choline. The pulse chase intravenous infusion experiment was an 18 h pulse of [trimethyl-^2^H_9_]choline followed by a 6 h chase during which nothing was infused. This experiment enables detection of choline that is derived from lipids. (e) Fraction of circulating choline derived from lipolysis. Values represent the labeled fraction of serum choline at the 24 h time point of the infusion normalized to the serum choline m+9 during the pulse. m+9 represents choline that has remained intact and comes from PC made in the Kennedy pathway. m+3 represents choline derived from PCs that were synthesized by the PEMT pathway. (f) Fraction of serum and tissue choline that is m+9 at the end of the chase experiment. The red line represents the fraction of serum choline that is labeled. Labeling below this line indicates serum choline as a potential source of labeling rather than tissue autonomous production. Tissues with more labeling than serum choline are likely sources of lipolysis-derived choline in circulation. Bars represent mean ± S.D. Each data point represents data from an independent mouse. P-values were calculated with ordinary one-way ANOVA with Dunnett correction for multiple comparisons. All experiments were performed in male *ad lib* fed C57Bl/6N mice during the light cycle. PC, phosphatidylcholine; PEMT, phosphatidylethanolamine methyltransferase; BHMT, betaine homocysteine methyltransferase

### Phosphatidylcholine is a major source of circulating choline

Choline is also a critical head group for phosphatidylcholine (PC) synthesis, a significant component of organelle, plasma, and lipoprotein membranes. PC can be synthesized directly from choline by the Kennedy pathway or produced by a series of three methylation reactions by PEMT in the liver. Because the PEMT pathway is the only pathway for the synthesis of choline, we were interested in how much choline in serum and in tissue comes from PC lipolysis. We can trace choline into PCs using [trimethyl-^2^H_9_]choline infusion, which results in M+9 PC if the Kennedy pathway is used and M+3 PC if choline contributes methyl groups to PEMT (Fig. 3a).

To measure how much circulating choline comes from these different PC pools, we performed pulse-chase [trimethyl-^2^H_9_]choline infusions in which we infused labeled choline for 18 hours to label PCs followed by a 6-hours chase during which time we stop the infusion to allow circulating metabolite labeling to decrease (Fig. 3b). If labeling of free choline is observed after this 6 hour chase period, it suggests choline production from PC lipolysis (Fig. 3b). After the 24 hour long experiment, we collected tissues and analyzed labeling of polar metabolites and lipids in serum and tissues. At the end of the 24-hour pulse-chase infusion, we observed broad labeling of PCs across tissues, demonstrating choline use for PC synthesis in the Kennedy pathway, which was consistent with our previous tracing of choline into tissue phosphocholine pools (Extended data Fig 2h). As expected, we observed M+3 labeling of PCs only in the liver, indicating contribution of PEMT to PC production (Fig. 3c). Labeling of PCs in the liver also showed a bias toward Kennedy pathway production for smaller, more saturated PCs and vice versa for larger, more unsaturated PCs (Extended data Fig 4a). Some tissue PC pools were labeled as much as the serum during the pulse (Fig. 3d), like white adipose and kidney, which suggests total turnover of the PC pools during the 24-hour experiment. Other tissues like the brain and fast muscle exhibited low labeling, suggesting slow turnover of their PC pool (Fig. 3c).

**Figure 4.**
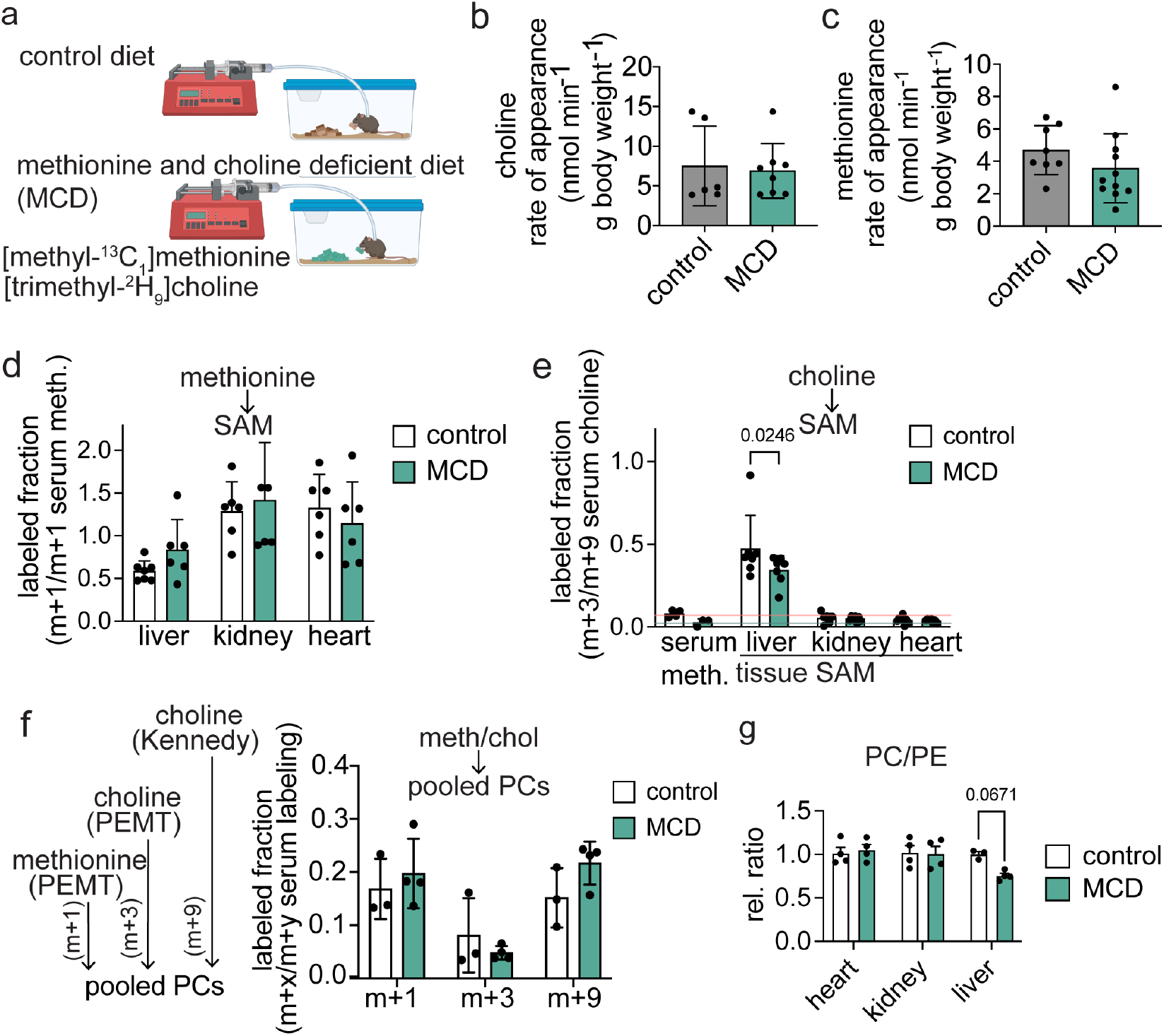
Systemic choline and methionine fluxes are maintained in animals fed methionine and choline deficient diet. (a) Schematic showing intravenous infusion of [trimethyl-^2^H_9_]choline and [methyl-^13^C_1_]methionine in mice fed a methionine and choline deficient diet or fed a paired control diet. (b) Rate of appearance of choline calculated from m+9 serum enrichments from 8 h intravenous infusion of [trimethyl-^2^H_9_]choline. (c) Rate of appearance of methionine calculated from m+1 serum enrichments from 8 h intravenous infusion of [methyl-^13^C_1_]methionine. (d) Normalized labeling of tissue SAM from 8 hour intravenous infusion of [methyl-^13^C_1_]methionine. (e) Normalized labeling of tissue SAM from 8 hour intravenous infusion of [trimethyl-^2^H_9_]choline. The red line represents the fraction of serum methionine that is labeled in control conditions and the green line represents the fraction of serum methionine that is labeled in MCD diet. Labeling below these lines indicates that labeling is potentially sourced from serum methionine pools rather than tissue autonomous production. (f) Normalized labeling of pooled PCs in the liver from 8 hour intravenous infusion of [trimethyl-^2^H_9_]choline. (g) Ratio of ion counts of pooled PC species to pooled PE species. Bars represent mean ± S.D. Each data point represents data from an independent mouse. P-values were calculated with two-way ANOVA with Šídák correction for multiple comparisons. All experiments were performed in male *ad lib* fed C57Bl/6N mice during the light cycle. PC, phosphatidylcholine; PE, phosphatidylethanoline; PEMT, phosphatidylethanolamine methyltransferase

In the serum, during the pulse intravenous infusion of [trimethyl-^2^H_9_]choline, about 14% of circulating choline was M+9 with a small fraction of PEMT-derived M+3 over time (Figure 3b). During the chase period, circulating choline labeling was about 6.8% and 1.8% M+9 and M+3, respectively. This suggests a large fraction of circulating choline is derived from PC lipolysis. To quantify the contribution of PC lipolysis, we divided the labeling observed at the end of the chase by the average labeling during the pulse, which approximates the maximum labeling of systemic PC pools. Based on this calculation, the PEMT pathway accounted for a minority of choline, accounting for only about 10% of the circulating choline (Figure 3e). In contrast, choline derived from lipolysis of PCs generated by the Kennedy pathway accounted for about 67% of circulating choline. These data highlight the significance of lipid recycling in supplying choline for body functions.

We next investigated in which tissue(s) PC lipolysis occurs. If we observe labeling of choline pools in tissue that is significantly higher than circulation, it would suggest internal tissue autonomous lipolysis. We observed M+9 choline labeling in the heart and gonadal white adipose tissue (gWAT) that was significantly higher than circulating choline at the end of the chase, suggesting significant hydrolysis of internal PC pools in these tissues (Fig. 3f). PCs in these tissues did show significant labeling but less than expected based on choline labeling, suggesting that there may be sub-pools of PCs that turnover faster than others (Extended data Fig. 4b). As expected, only the liver showed a higher fraction of M+3 choline compared to serum was the liver (Extended data Fig. 4c). Thus, we find that the majority of choline comes from PC hydrolysis with a minority coming from PCs made by de novo synthesis through PEMT.

### Systemic choline and methionine fluxes are maintained despite changes in liver methyl group fate

Because we observed a significant contribution of choline to the hepatic catabolic pathway, we sought to understand how choline fate is altered in response to dietary methyl deficiency. To do this, we used a methionine and choline-deficient (MCD) diet with a matched control diet. MCD diets are known to lead to increased hepatic steatosis and fibrosis but lack the metabolic phenotypes of MASLD, such as insulin resistance or weight gain. Instead, animals are known to become hypermetabolic ^22^. Our goal in using this diet is to understand how the body responds to choline deficiency, not to model MASLD. Thus, we put animals on a 3-week diet to model dietary methyl-donor deficiency, but not liver disease. To investigate how methionine and choline metabolism is altered under these dietary conditions, we co-infused [methyl-^13^C_1_]methionine and [trimethyl-^2^H_9_]choline (Fig 4a). Surprisingly, choline and methionine rates of appearance were unchanged despite deficiency of these nutrients in the diet (Fig 4b-c). In addition, contributions of choline and methionine to SAM synthesis were only minimally altered across all measured tissues, with a significant difference in the contribution of choline to SAM driven by a single outlier (Fig 4d-e). SAM levels were normal across all tissues but the liver, despite decreased methionine in the heart, kidney, and pancreas (Extended data Fig. 5a-b). Choline levels were only impacted in the liver (Extended data Fig 5c). These data suggest that despite a severe dietary deficiency, systemically circulating choline fluxes, circulating methionine fluxes, and the contribution of each to the methionine cycle flux are largely maintained.

**Figure 5.**
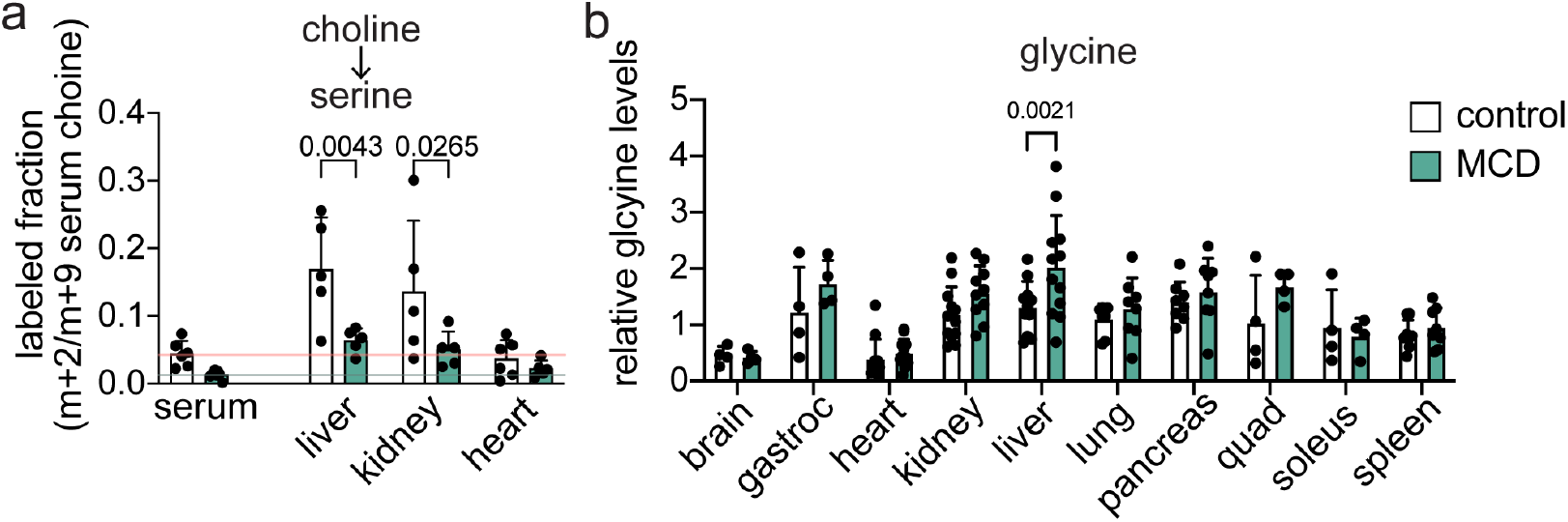
Methionine and choline deficient diet reduces the contribution of choline to the mitochondrial folate cycle. (a) Normalized labeling of serum and tissue serine from 8 hour intravenous infusion of [trimethyl-^2^H_9_]choline.The red line represents the fraction of serum serine that is labeled in control conditions and the green line represents the fraction of serum serine that is labeled in MCD diet. Labeling below these lines indicates that labeling is potentially sourced from serum serine pools rather than tissue autonomous production. (b) Relative ion counts for glycine across tissues in MCD diet and control diet fed mice. Bars represent mean ± S.D. Each data point represents data from an independent mouse. P-values were calculated with two-way ANOVA with Šídák correction for multiple comparisons (a) or Mixed-effects analysis with Šídák correction for multiple comparisons (b). All experiments were performed in male *ad lib* fed C57Bl/6N mice during the light cycle. MCD, methionine and choline deficient

To further investigate the impact of the MCD diet on the tissue methionine cycle, we next analyzed production of methyltransferase reactions. Because SAM and choline were decreased in the liver, we investigated PC synthesis pathways. Surprisingly, labeling of PCs through both the PEMT and the Kennedy pathway was maintained in the liver (Fig. 4f). PC levels were also maintained, indicating maintained PC synthesis flux (Extended Data Fig. 5D). However, the PC/PE ratio trended down in the MCD diet, suggesting a modest accumulation of PE (Fig 4g).

Other high-flux methlytransferases include glycine n-methyltransferase (GNMT), which produces sarcosine from glycine and guanidinoacetic acid n-methyltransferase (GAMT), which produces creatine from guanidinoacetic acid. GNMT is highly expressed in the liver and pancreas, and its main role is thought to be to prevent SAM from becoming too high in the cytosol^23,24^. Levels of sarcosine were only changed in the liver, suggesting that in the MCD diet, there is less oversupply of SAM for GNMT to alleviate (Extended data Fig. 6a). Combined with our data that supports sustained PC synthesis flux with MCD diet, this data argues that choline deficiency causes SAM to be prioritized for PC synthesis. To gain a deeper understanding of the competition for SAM in the liver, we compared published values for enzyme kinetics of PEMT and GNMT in the Brenda database. We found that PEMT has a lower K_m_ than GNMT (Extended data Fig. 6b). This suggests that decreased SAM levels may limit its use by GNMT. In line with the competition for SAM between PEMT and GNMT, previous studies have shown that GNMT knockout increases PC synthesis^23^. GAMT is expressed in more tissues including the liver, pancreas, muscle and kidney, but we only saw decreased levels of creatine in the pancreas (Extended data Fig. 6c). Corresponding to changes in methylation potential in the liver and pancreas, we saw accumulation of the substrate of GAMT, guanidinoacetic acid, in both tissues (Extended data Fig. 6d). Overall, despite dietary methionine and choline deficiency, tissue methionine cycle is broadly maintained with sustained PC synthesis flux and small changes in methylation potential in the liver and pancreas.

**Figure 6.**
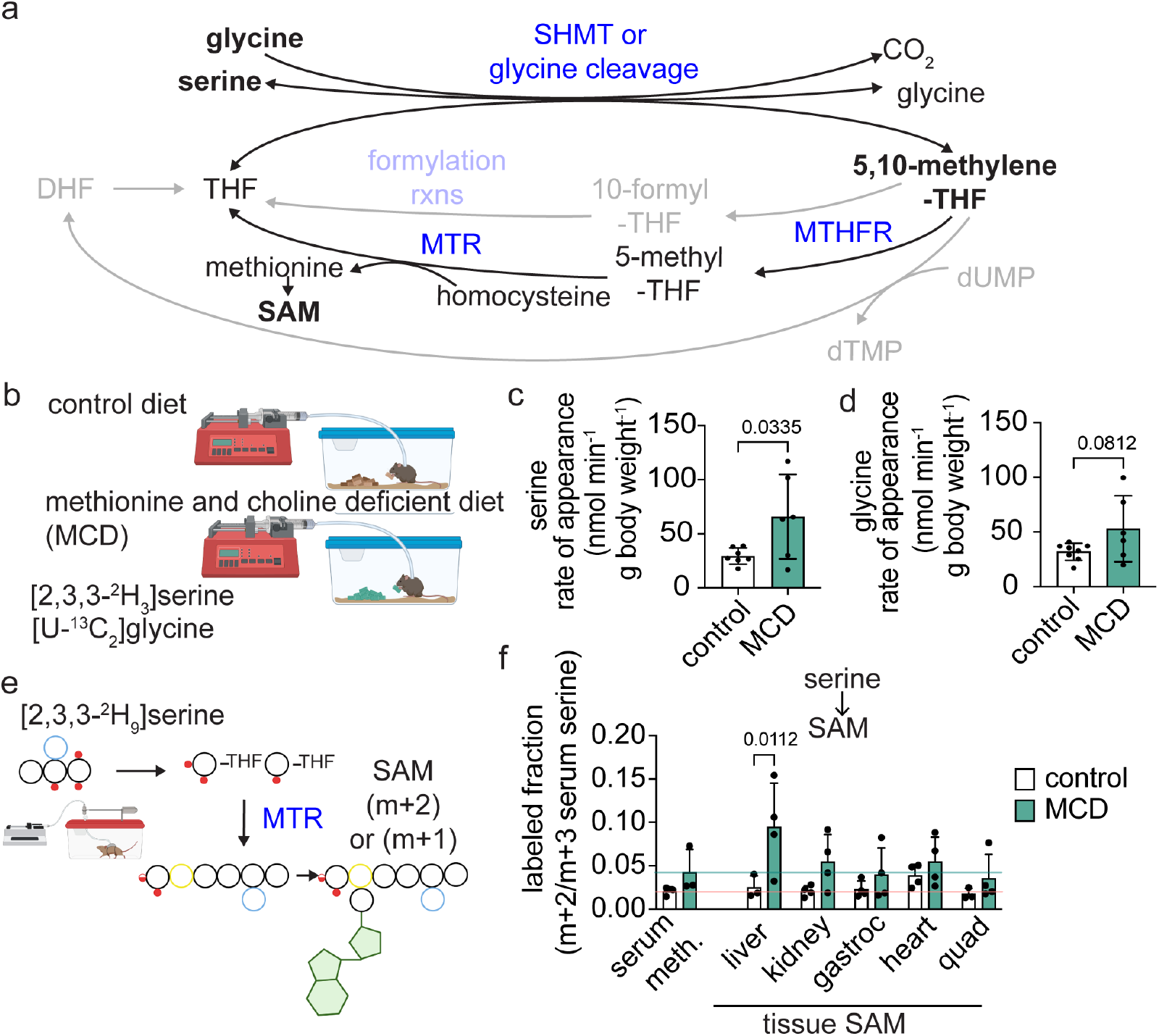
Dietary methionine and choline deficiency triggers increased systemic serine flux to support SAM synthesis in the liver. (a) Schematic of the folate cycle highlighting reactions that support production of 5,10-methylene-THF and SAM. (b) Schematic showing intravenous infusion of [2,3,3-^2^H_3_]serine and [U-^13^C_2_]glycine in mice fed a methionine and choline deficient diet or fed a paired control diet. (c) Rate of appearance of serine calculated from m+3 serum enrichments from 8 h intravenous infusion of [2,3,3-^2^H_3_]serine. (d) Rate of appearance of glycine calculated from m+2 serum enrichments from 8 h intravenous infusion of [U-^13^C_2_]glycine. (e) Schematic of use of [2,3,3-^2^H_3_]serine tracing to measure the contribution of the folate cycle to SAM synthesis. (f) Normalized labeling of tissue SAM from 8 hour intravenous infusion of [2,3,3-^2^H_3_]serine. The red line represents the fraction of serum methionine that is labeled in control conditions and the green line represents the fraction of serum methionine that is labeled in MCD diet. Labeling below these lines indicates that labeling is potentially sourced from serum methionine pools rather than tissue autonomous production. Bars represent mean ± S.D. Each data point represents data from an independent mouse. P-values were calculated with two tailed T-test (c,d) or two-way ANOVA with Šídák correction for multiple comparisons (f). All experiments were performed in male *ad lib* fed C57Bl/6N mice during the light cycle. MCD, methionine and choline deficient; SHMT, serine hydroxymethyltransferase, MTR, 5-methyltetrahyrofolate homocysteine methyltransferase; SAM, s-adenosylmethionine

### Methyl-unit deficiency diminishes the contribution of choline to the mitochondrial folate cycle

Because we saw changes in liver methylation fate, we analyzed choline catabolism further downstream of the methionine cycle at the DMGDH and SARDH steps. Like in Figure 2, we used data from our [trimethyl-^2^H_9_]choline infusion to look at the contribution of choline-derived methyl groups into the mitochondrial folate cycle. Because serine is net produced in the liver, which consumes 5,10-methylene-THF, we again used serine labeling as a readout of contribution to the mitochondrial folate cycle. Contribution of [trimethyl-^2^H_9_]choline to serine was considerably decreased in both the liver and kidney, suggesting that the contribution of choline-derived dimethylglycine and sarcosine to the mitochondrial folate cycle is impaired (Fig. 5a). To confirm that one carbon units are used in the liver and kidney for serine synthesis in the context of MCD diet, we infused [U-^13^C_2_]glycine (Extended data Fig. 7a). Serine labeling from glycine was unchanged in the liver and kidney, indicating that serine synthesis is still occurring. Consistent with a shortage of one-carbon units in the mitochondria of the liver, we also saw increased glycine levels with MCD diet only in the liver (Fig 5b). Thus, these data suggest that choline deficiency leads to depletion of one-carbon units in liver mitochondria.

**Figure 7.**
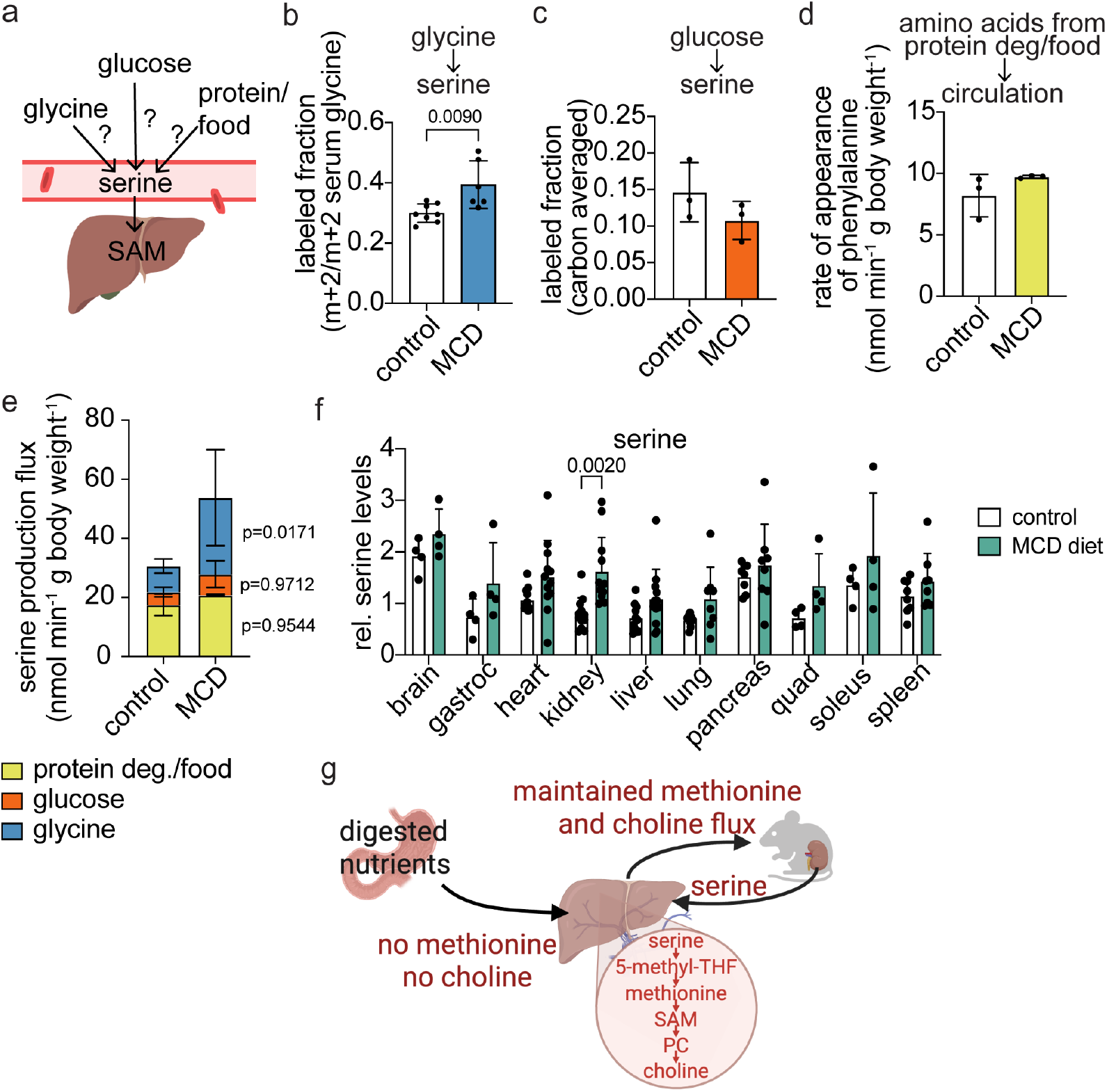
Increased systemic serine flux is supported by increased production from glycine. (a) Schematic showing potential sources of circulating serine. (b) Normalized labeled fraction of serum serine from infusion of [U-^13^C_2_]glycine. (c) Normalized labeled fraction of serum serine from infusion of [U-^13^C_6_]glucose. (d) Rate of appearance of essential amino acid phenylalanine from intravenous infusion of [^15^N]phenylalanine. (e) Serine production fluxes from different sources measured using the serine rate of appearance and infusions of U-^13^C_2_]glycine, U-^13^C_6_]glucose, and [^15^N]phenylalanine. (f) Relative ion counts for serine across tissues in MCD diet and control diet fed mice. (g) Schematic of proposed mechanism. When methionine and choline are deficient in the diet, there is increased release of serine from the kidney that supports maintained methionine, PC, and choline synthesis in the liver. Thus, methionine and choline fluxes are maintained by one-carbon metabolism buffering. Bars represent mean ± S.D. Each data point represents data from an independent mouse. P-values were calculated with two tailed T-test (b), two-way ANOVA with Šídák correction for multiple comparisons (e), or mixed-effects analysis with Šídák correction for multiple comparisons (f). All experiments were performed in male *ad lib* fed C57Bl/6N mice during the light cycle. SAM, s-adenosylmethionine; MCD, methionine and choline deficient

### Increased systemic serine and glycine fluxes compensate for methyl-donor deficiency

To this point, we have observed that dietary methionine and choline deficiency lead to small changes in SAM and methyl group fate and decreased production of 5,10-methylene-THF in liver mitochondria. Synthesis of both SAM and 5,10-methylene-THF can be supported by the input of nutrients such as serine and glycine (Figure 6a). Thus, if production from choline is decreased, we hypothesized that other one-carbon donors, such as serine or glycine, must feed in to compensate. One carbon from serine can be transferred to THF by serine hydroxymethyltransferase, producing glycine and 5,10-methylene-THF. Glycine cleavage can also produce a 5,10-methylene-THF from THF and a CO_2_. One-carbon units carried by THF in the form of 5,10-methylene-THF can then either be used directly to make dTMP from dUMP or converted to 10-formyl-THF for various formylation reactions or 5-methyl-THF for methylation of homocysteine to make methionine (Fig. 6a).

To determine if one-carbon units from serine or glycine contribute to the methionine cycle, we independently intravenously infused [2,3,3-^2^H_3_]serine and further analyzed our data from [U-^13^C_2_]glycine infusions in MCD and control diet conditions (Fig 6b). Strikingly, serine rate of appearance was increased substantially in MCD conditions with the glycine rate of appearance also trending up (Fig 6c-d). This suggests that in response to dietary deficiency of methionine and choline, one or more tissues increase the release of serine into circulation. The change in rate of appearance of serine was much larger than what was previously observed with serine and glycine-deficient and sufficient diets^19^, suggesting a major rewiring of folate cycle metabolism.

While we were surprised by the extent of the systemic increase in serine turnover, it is in line with our hypothesis that serine or glycine’s contribution to 5,10-methylene-THF and methionine synthesis is increased with feeding of MCD diet (Fig 6e). Therefore, we looked at labeling of SAM in tissues (Fig 6f and Extended data 7b) and found that in MCD conditions, there was a significant increase in labeling of SAM only in the liver (Fig 6f). Labeling from glycine into SAM pools was inconsistent and low (Extended data Fig. 7b), suggesting that most one-carbon units used to compensate for methionine and choline deficiency come from serine. These data suggest that in response to methionine and choline deficiency, the liver uses serine-derived one-carbon units from the folate cycle to support methionine synthesis via 5-methyltetrahydrofolate homocysteine methyltransferase (MTR).

While it has been suggested that increased serine use in the methionine cycle could compensate for methionine and choline deficiency previously^25^, the large increase in serine rate of appearance that we observed suggests an unexpected systemic remodeling of metabolism, which supports an expected outcome of hepatic methionine production. The about two-fold increase in serine rate of appearance indicates that there must be increased synthesis and release of serine from one or more tissues. Therefore, we next investigated what metabolic pathways and which tissue supports increased serine rate of appearance in methionine and choline-deficient diet-fed animals.

### Increased serine production from glycine supports increased rate of appearance

Serine in the bloodstream can come from diet, protein degradation, or through 2 synthesis pathways from other nutrients: 1-Production of serine can occur from glycine through reverse flux of serine hydroxymethyltransferase (SHMT), which requires a one-carbon unit from 5,10-methylene-THF and 2-from glucose-derived 3-phosphoglycerate, which can be used in the *de novo* serine synthesis pathway with a nitrogen from glutamate (Fig. 7a). To distinguish between these different sources of serine, we performed independent stable isotope tracing experiments to analyze each of these pathways.

To look at serine synthesis from glycine, we analyzed our data from the [U-^13^C_2_]glycine infusion experiment. Labeling of serine from glycine in serum was significantly increased with MCD diet (Fig 7b), while labeling of glycine from the [2,3,3-^2^H_3_]serine infusion was unchanged (Extended data Fig 8). This suggests increased net serine production from glycine via reverse SHMT, but not vice versa. To look at the *de novo* serine synthesis pathway, we intravenously infused [U-^13^C_6_]glucose. To account for serine from protein degradation and dietary serine, we co-infused isotope-labeled glucose with essential amino acid [^15^N]phenylalanine and measured its rate of appearance. Because essential amino acid phenylalanine can only come from diet or protein degradation, if one of these is changed, we should observe a change in phenylalanine’s rate of appearance. However, there was no change in the labeling of serum serine from glucose (Fig 7c) or the rate of appearance of phenylalanine in methionine and choline-deficient conditions (Fig 6d). Combined, these data suggest that increased serine synthesis is supported by increased net production of serine from glycine.

To further confirm the source of serine that enables an increased rate of appearance of serine in MCD diet conditions, we calculated serine production flux values from our infusion experiment data. We calculated serine production flux from protein degradation and diet using the phenylalanine rate of appearance by adjusting for the frequency of serine in protein relative to phenylalanine, similar to what was done previously^19^. There was no difference in serine production from these sources between the control and MCD diet (Fig. 7e). To calculate serine synthesis flux from glycine and from glucose, we used the labeling data from [U-^13^C_6_]glucose and U-^13^C_2_]glycine infusions (Fig 7b-c) and the serine rate of appearance (Fig 6d). This calculation revealed a significant increase in the serine production flux from glycine that accounted for the majority of the increase in serine rate of appearance in MCD diet (Fig 7e).

Next, we aimed to identify which tissue or tissues were responsible for increased serine production. As mentioned previously, tissue labeling did not reveal any significant labeling of serine from glycine at the tissue levels, but a trend toward increased labeling was observed in the kidney (Extended data Fig 7a). In addition, we observed increased serine levels in the kidney (Fig 7f). Combined, these data suggest that there is remodeling of the folate cycle to support serine production in the kidney and to increase serine rate of appearance. This increased serine release enables greater methionine synthesis in the liver by providing one-carbon units for the folate cycle, which supplies 5-methyl-THF for use by MTR (Fig. 7g). This supports a model in which the liver can compensate for dietary deficiencies of methionine and choline. To do this, serine is produced by the kidney to support hepatic methionine synthesis from folate cycle-derived methyl groups.

## Discussion

Insufficient consumption of choline is common in the human population^5^, yet we have a limited understanding of how remodeling of systemic metabolism in response to this insufficiency could contribute to disease development. In this study, we investigate how choline is used in the body and find that choline catabolism is uniquely important in supporting one-carbon metabolism in the liver. To gain a broader understanding of how choline deficiency or insufficiency may impact one-carbon metabolism and tissue function, we investigated how MCD diet influences one-carbon fluxes. While excluding methionine and choline from the diet is extreme, it serves as a clean model of choline deficiency to reveal the pathways impacted. Surprisingly, we found that systemic methionine and choline fluxes are maintained after 3 weeks on the MCD diet. However, we found two other metabolic changes in response to dietary methyl donor deficiency. We found that the choline-derived methyl groups no longer contributed to the mitochondrial folate cycle in the liver. In addition, we uncovered changes in systemic serine metabolism that compensate for dietary methionine and choline deficiency by enabling methionine synthesis in the liver.

Notably, the liver bears the majority of the burden in response to dietary methyl deficiency, losing a major one-carbon source and compensating by increasing methionine synthase activity and maintaining PC production. Decreased PC levels have been considered the main mechanism of hepatic steatosis in the MCD diet by impairing the liver’s ability to export fat via VLDL particles, which have lipid monolayer membranes enriched in PCs relative to the ER membrane. Our data supports maintained PC synthesis and maintained PC levels in early choline deficiency by redirecting SAM from sarcosine production via GNMT to sustain PEMT flux. However, we still observed a decrease in the PC/PE ratio, consistent with the idea that small changes in PC/PE may influence VLDL export rates.

This study uncovered choline’s contribution to the liver mitochondrial folate cycle, which was deficient in MCD diet conditions. Human variants of the first enzyme in this pathway DMGDH are associated with the lipid composition of lipoproteins^26–29^. DMGDH and the downstream enzyme SARDH support mitochondrial function by supplying one-carbon units into the folate cycle to produce nucleotides and formylmethionine-tRNA, which supports mitochondrial DNA replication, transcription, and translation. These reactions also donate electrons into the ETC via the coenzyme Q pool^21^. How DMGDH may influence lipoprotein composition is unclear, but decreased mitochondrial function may relate to changes in fatty acid oxidation or VLDL secretion, as VLDLs are produced at ER-mitochondria contact sites^30^. Combined with human genetic data, our study highlights the potential influence that this pathway may have on the development of hepatic steatosis.

We also found systemic increases in fluxes of nutrients that feed the folate cycle in response to MCD diet. The folate cycle plays an essential role in human health and disease development^31^, and folate cycle-derived one-carbon units support nucleotide synthesis and are thus implicated in cancer^15,32^. Choline consumption is correlated with reduced cancer risk^33,34^, and thus increased serine flux in the context of choline deficiency provides a potential link between choline and folate-cycle-supported nucleotide synthesis to could support proliferation in the context of cancer. Our data also supports increased serine synthesis in the kidney, which suggests changes in kidney one carbon metabolism. Dietary choline consumption has also been anticorrelated with chronic kidney disease^35^, and our study provides a link between choline deficiency and altered kidney metabolism. Thus, our study provides novel metabolic mechanisms of how systemic metabolism can buffer for dietary deficiency of choline and reveals tissue crosstalk to coordinate this compensatory response.

## Methods

### Mouse experiments

Mouse work was approved by the University of California Los Angeles Institute Animal Care and Use Committee (protocol no. 2022-119). Mice were housed in a temperature- and humidity-controlled vivarium on a 12-hour light/dark cycle with ad libitum access to food and water. Standard chow was provided unless otherwise noted (PicoLab Rodent 20 5053). Eleven-week-old wild type C57Bl/6NCrl mice were purchased from Charles River Laboratories (strain code no. 027) and given at least 5 days to acclimate before experiments.

For intravenous infusion of stable isotope tracer, aseptic in-house catheterization of the right jugular vein was performed, and the catheter was connected to a vascular access button (Instech, VABM1B/25) for externalization. Following surgery, mice were individually housed until sacrifice and allowed one week recovery before any experiments were performed. Infusions were performed in the home cage using a swivel-tether system (Instech) that allows free movement around the cage while maintaining regular access to food and water gel cups (Bio-Serv or ClearH2O). All tracers were prepared in a 0.9% saline solution, with details on tracer concentrations and infusion rates provided in Table 1. Unless overwise noted, infusions were 8 hours starting at approximately 10 am and ending at 6pm. Tail blood was collected at defined intervals using blood collection tubes with clotting factor (Starstedt 16.442.100) to analyze circulating metabolite labeling. Blood samples were kept on ice to promote coagulation, then centrifuged for 5 minutes at 4 °C to isolate serum. Tissue samples were snap-frozen with a liquid nitrogen-cooled Wollenberger clamp and stored on dry ice or at −80 °C to ensure preservation of metabolite integrity prior to analysis.

### Dietary intervention

Methyl deficiency was induced by transitioning mice from standard chow to a methionine and choline-deficient (MCD) diet (MP Biomedicals, MP296043910), while age-matched controls received a matched methionine and choline replete control diet (MP Biomedicals, MP296044110). Diet interventions began on day 0, and mice were maintained on their assigned diets for three weeks.

### Metabolite extraction for LC-MS analysis

For tissue, polar metabolites were extracted from snap-frozen tissue samples using a standardized cryogenic grinding protocol followed by liquid-liquid extraction. Precooled 2 mL Eppendorf microcentrifuge tubes containing grinding beads were utilized for cryogenic milling at 25 rps for 30–60 seconds to achieve thorough tissue pulverization. The resulting tissue powder was transferred to precooled tubes, weighed, and extracted with a 40:40:20 (v/v/v) methanol:acetonitrile:water solution (40 µL per mg of tissue). Samples were vortexed and incubated on dry ice for 30 minutes or stored at −80°C overnight to enhance metabolite solubilization. Following centrifugation at 4°C at maximum speed for 30 minutes, the supernatant was carefully transferred to 1.5 mL Eppendorf tubes and stored at −80°C until analysis. Prior to LC-MS injection, samples underwent an additional 10-minute centrifugation step, and 120 µL of the clarified supernatant was transferred to an MS vial for metabolomic analysis.

For extraction from sera, methanol (100%) was precooled on dry ice, while serum samples were thawed on wet ice. 5 µL of serum was transferred into each microcentrifuge tube, followed by the addition of 145 µL of 100% methanol. The samples were vortexed thoroughly to ensure complete homogenization. The resulting mixtures were incubated on dry ice for 30 minutes or frozen at −80°C overnight. Following incubation, the samples were centrifuged at maximum speed (4°C) for 30 minutes. After centrifugation, 100 µL of the supernatant was carefully transferred into new microcentrifuge tubes. A second centrifugation step was performed at maximum speed (4°C) for 10 minutes. Finally, 75 µL of the resulting supernatant was transferred to MS vials for analysis.

### Polar metabolite LC-MS analysis

Polar metabolites were separated on a Waters BEH z-HILIC column (150 × 2.1 mm, 2.7 µm) using a Thermo Scientific™ Orbitrap Exploris™ 480 mass spectrometer with H-ESI source. Column and autosampler temperatures were maintained at 25°C and 4°C, respectively. The mobile phase comprised 10 mM ammonium bicarbonate in water (Solvent A, pH ∼9.1) and 95% acetonitrile/5% water (Solvent B). Chromatography was performed at 0.18 mL/min with a gradient from 95% to 45% B over 15 min, held at 45% B, ramped back to 95% B, and re-equilibrated. MS data were acquired in DDA mode with fast polarity switching. MS1 scans were performed at 120,000 resolution (m/z 70–1000; AGC target 1 × 10^6^). MS/MS scans used a TOP2 ddMS2 method with stepped HCD (10% and 40%), 30,000 resolution, and a 0.7 m/z isolation window. Dynamic exclusion, isotope exclusion, monoisotopic precursor selection, and mild ion trapping were enabled to optimize spectral quality. Raw LC-MS data were converted to mzXML (MSConvert) and analyzed in El MAVEN (v0.12.0) with default settings for peak detection, chromatogram alignment, and isotope annotation. Retention times were normalized to external standards, ion intensities corrected for matrix effects, and peaks manually curated when necessary. Processed data were exported as CSV files for analysis of isotopologue distributions.

### Non-polar metabolite LC-MS analysis

Non-polar metabolites were profiled by LC-MS using Thermo Scientific™ Orbitrap Exploris™ 480 mass spectrometer with H-ESI source operating in polarity switching mode. MS1 scans were collected at 120,000 resolution (200 m/z) over 300–1200 m/z. DDA selected the seven most abundant precursor ions for MS2, using 30,000 resolution, stepped HCD (10/40), a 1.5 m/z isolation window, 5 × 10^3^ intensity threshold, 20 s dynamic exclusion, and 30% apex trigger. Separation was performed with an Agilent InfinityLab Poroshell aQ-C18 column (150 mm × 2.1 mm, 2.7 µm) at 50°C, 0.27 mL/min flow, and 5 µL injection. The mobile phases were (A) water with 0.2% formic acid and 2 mM ammonium formate, and (B) methanol with 0.2% formic acid and 2 mM ammonium formate. The 29.46-minute gradient started at 70% B for 0.44 min, increased to 90% B over 3.61 min, to 99% B over 6.75 min, held for 9.9 min, reduced to 30% B over 0.45 min, and re-equilibrated at 30% B for 8.31 min. For pooled lipid analysis, ion counts for all lipids of a given species were added together. When pooling lipid labeling data, we summed the ion counts for each isotopologue for all lipid species of that class in which we detected labeling (e.g all PC M+0, all PC M+1, etc.). Then, for calculation of labeled fraction, we divided the sum of the isotopologues of interest (sum of all PC M+9) by the sum of all isotopologues of all lipids of that class (sum of all PC) in which we were able to detect labeling.

### Rate of appearance calculation

To measure the rate of appearance, alternatively termed the circulatory turnover flux^14^, we used our intravenous infusion of stable isotope tracers data. At pseudo-steady state, we measure the isotope distribution in serum and define the fraction of the metabolite in serum that is fully labeled after the tracer enrichment has reached its plateau as *L* (e.g. M+9 for [trimethyl-^2^H_9_]choline) and the rate of infusion of the tracer *R*, in units of nmol × min^-1^ × g bodyweight^-1^. Thus, the rate of appearance (*R*_*a*_) of a given traced metabolite was calculated given the equation 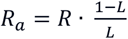

### Quantification and statistical analysis

Statistical analysis was performed using GraphPad Prism 9 software. Data with normal distribution are presented as means ± standard deviation (SD), unless stated otherwise. Statistical tests used for each data set are indicated in figure legends.

