## Supplementary material for "Flexibility of systemic one-carbon metabolism partially buffers dietary methyl donor deficiency": Table 1 and Extended data

**Contents:**

Table 1
Extended Data Figures 1-8

| Tracer | Concentration | Infusion rate ($\mu$l · min$^{-1}$ · g body weight$^{-1}$) | Infusion flux (nmol· min$^{-1}$ · g body weight$^{-1}$) |
|---|---|---|---|
| [methyl-$^{13}C_1$]methionine | 10 mM | 0.1 | 1 |
| [trimethyl-$^{2}H_9$]choline | 15 mM | 0.1 | 1.5 |
| [$^{13}C_2$]choline | 15 mM | 0.1 | 1.5 |
| [2,3,3-$^{2}H_3$]serine | 50 mM | 0.1 | 5 |
| [$^{13}C_2$]glycine | 40 mM | 0.1 | 4 |
| [U$^{13}C_6$]glucose | 400 mM | 0.1 | 40 |
| [$^{15}N_1$]phenylalanine | 20 mM | 0.1 | 2 |

**Table 1: Tracers used and infusion rate used in isotope tracing experiments.**

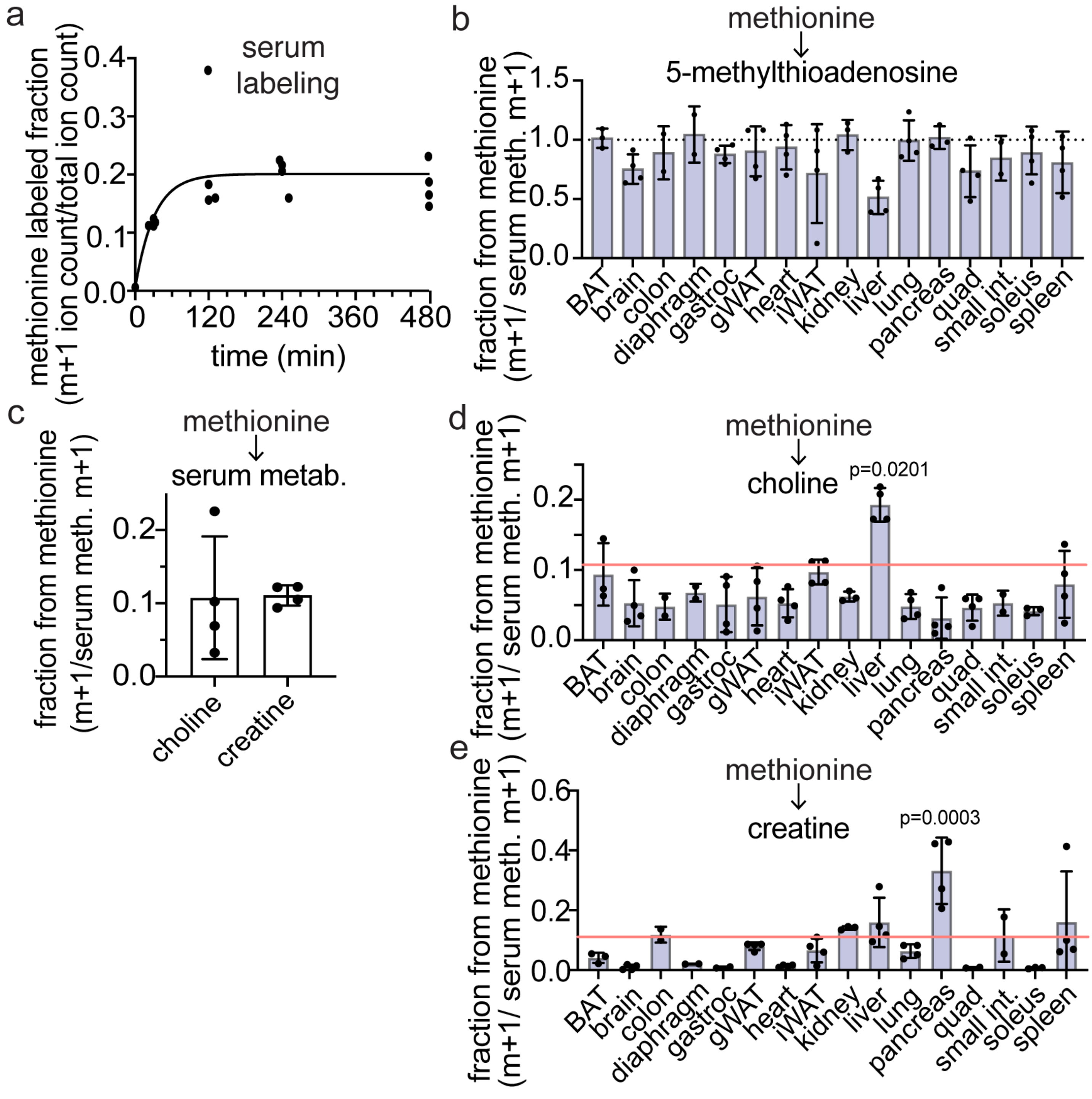


**Extended Data Figure 1. Methionine supplies methyl groups for all tissues.**

(a) Methionine labeling in serum over time during intravenous infusion of [methyl-$^{13}C_1$]methionine. Line represents one phase association curve fit to the data.

(b) Labeling of tissue 5-methylthioadenosine (MTA) normalized to serum enrichment after 8 hour intravenous infusion of [methyl-$^{13}C_1$]methionine representing the fraction of MTA derived from circulating methionine. The dotted line indicates a labeled fraction of 1, which would imply complete labeling of the tissue MTA pool from methionine.

(c) Labeling of serum m+1 choline and m+1 creatine normalized to serum enrichment after 8 hour intravenous infusion of [methyl-$^{13}C_1$]methionine representing the fraction of each circulating metabolite that is derived from circulating methionine.

(d) Labeling of tissue choline normalized to serum enrichment after 8 hour intravenous infusion of [methyl-$^{13}C_1$]methionine representing the fraction of choline derived from circulating methionine. Red line indicates labeling observed in serum choline. Labeling in the liver was significantly higher than serum suggesting internal production of choline from methionine.

(e) Labeling of tissue creatine normalized to serum enrichment after 8 hour intravenous infusion of [methyl-$^{13}C_1$]methionine representing the fraction of creatine derived from circulating methionine. Red line indicates labeling observed in serum creatine. Labeling in the pancreas was significantly higher than serum suggesting internal production of creatine from methionine.

Bars represent mean $\pm$ S.D. P-values were calculated with one-way ANOVA with Dunnett correction for multiple comparisons. Each data point represents data from an independent mouse. All experiments were performed in male *ad lib* fed C57Bl/6N mice during the light cycle.

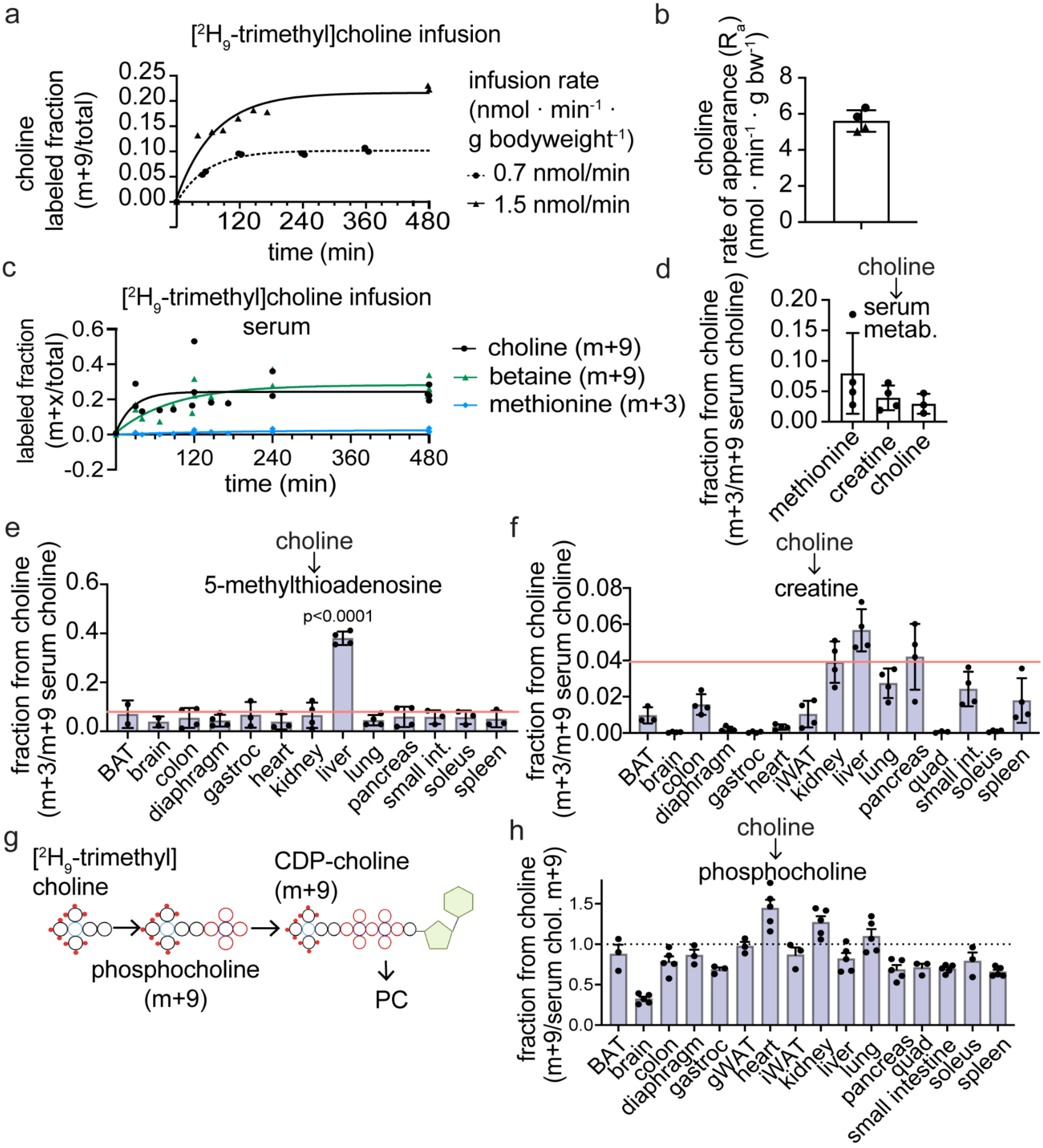


**Extended Data Figure 2. Choline is used in all tissues, and catabolism is active in the liver.**

(a) Labeled fraction of choline in in serum over time during intravenous infusion of [trimethyl-$^{2}H_{9}$]choline when infused at 0.7 (circles) or 1.5 (triangles) nmol · $min^{-1}$ · g $bodyweight^{-1}$. Lines represent one phase association curve fit to the data.

(b) Choline rate of appearance calculated from serum enrichments observed during intravenous infusion of [trimethyl-$^{2}H_{9}$]choline in (a). Values were calculated from n=2 infusions at 0.7 nmol · $min^{-1}$ · g $bodyweight^{-1}$ (circles) and n=2 infusions at 1.5 nmol · $min^{-1}$ · g $bodyweight^{-1}$ (triangles).

(c) Labeled fraction of choline, betaine, and methionine in serum over time during intravenous infusion of [trimethyl-$^{2}H_{9}$]choline. Choline data from 2 mice are replotted from panel a. Lines represent one phase association curve fit to the data.

(d) Labeling of serum metabolites methionine, creatine, and choline normalized to serum enrichment of choline after 8 hour intravenous infusion of [trimethyl-$^{2}H_{9}$]choline representing the fraction of each metabolite that comes from methylation reactions powered by choline-derived methyl units.

(e) Labeling of tissue 5-methylthioadenosine (MTA) normalized to serum enrichment of choline after 8 hour intravenous infusion of [trimethyl-$^{2}H_{9}$]choline representing the fraction of MTA that comes from choline.

(f) Labeling of tissue creatine normalized to serum enrichment of choline after 8 hour intravenous infusion of [trimethyl-$^{2}H_{9}$]choline representing the fraction of creatine that comes from choline.

(g) Schematic showing tracing of [trimethyl-$^{2}H_{9}$]choline into the Kennedy pathway.

(h) Labeling of tissue phosphocholine normalized to serum enrichment of choline after 8 hour intravenous infusion of [trimethyl-$^{2}H_{9}$]choline representing the fraction of phosphocholine that comes from choline.

Bars represent mean $\pm$ S.D. P-value was calculated with one-way ANOVA with Dunnett correction for multiple comparisons. Each data point represents data from an independent mouse. All experiments were performed in male *ad lib* fed C57Bl/6N mice during the light cycle.

Kapelczak, et al., Extended data Fig 3

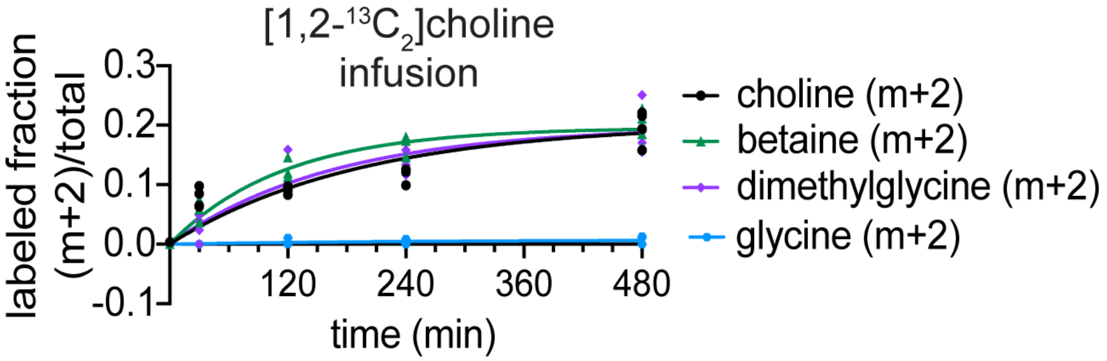


**Extended Data Figure 3. Choline infusion results in complete labeling of betaine and dimethylglycine pools, but not glycine.** Labeled fraction of choline in serum over time during intravenous infusion of [1,2-$^{13}C_2$]choline. Lines represent one phase association curve fit to the data. Each data point represents data from an independent mouse. All experiments were performed in male *ad lib* fed C57Bl/6N mice during the light cycle.

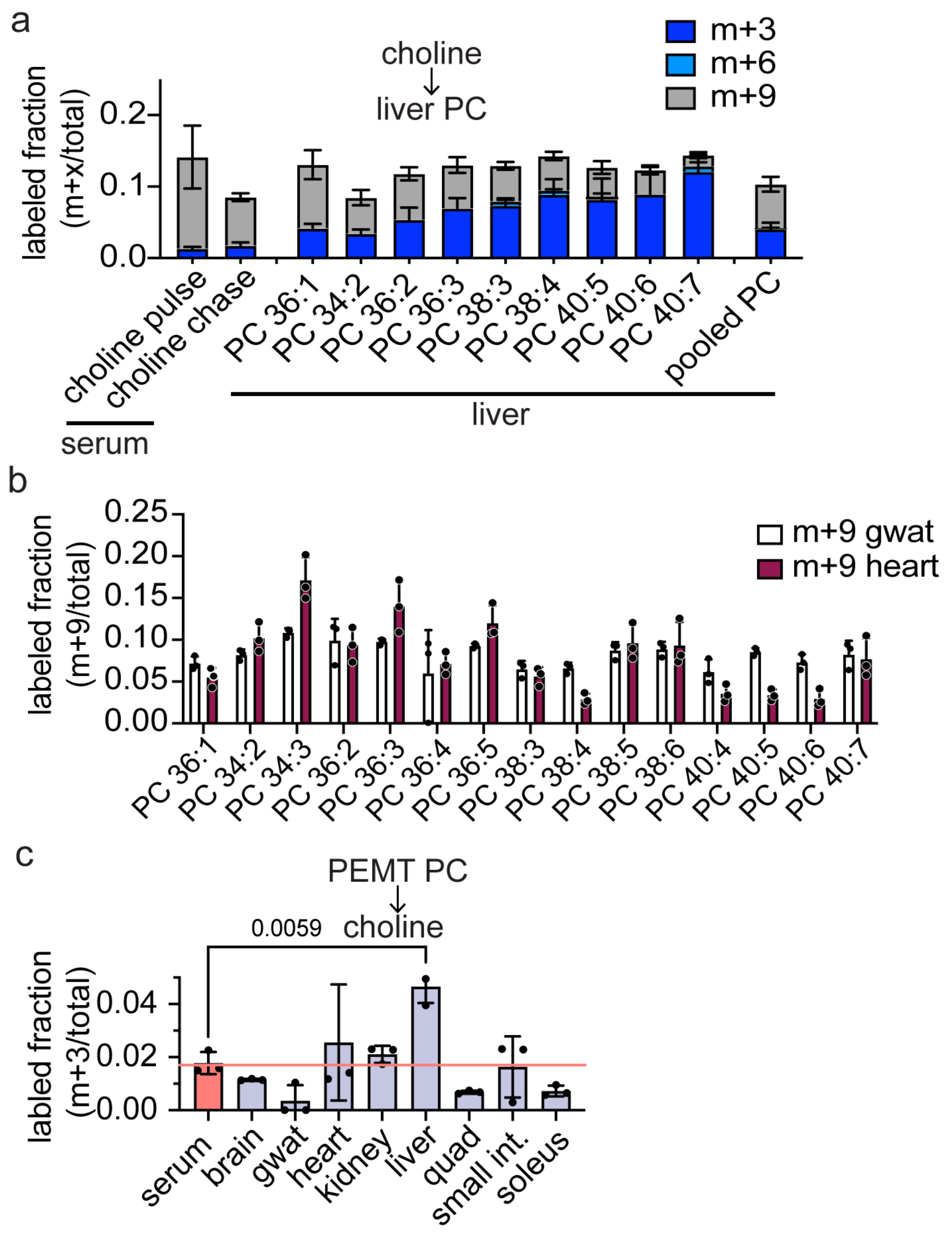


**Extended Data Figure 4. Labeling of choline species after pulse chase infusion of [trimethyl-$^{2}H_{9}$]choline.**

(a) Fraction of serum choline that is labeled at the end of the 18 h pulse infusion of [trimethyl-$^{2}H_{9}$]choline, serum choline after the 6 hour chase, and tissue phosphatidylcholine in liver that is labeled at the end of the 24 h pulse chase experiment.

(b) Fraction of gonadal white adipose tissue (gwat) and heart PC that is labeled at the end of the [trimethyl-$^2H_9$]choline pulse chase experiment.

(c) Fraction of serum and tissue choline that is labeled at the end of the [trimethyl-$^2H_9$]choline pulse chase experiment. Red line represents the labeling of serum choline, and labeling at or below this line suggests uptake of labeled choline rather than autonomous production from lipolysis.

Bars represent mean $\pm$ S.D. P-value was calculated with one-way ANOVA with Dunnett correction for multiple comparisons. Each data point represents data from an independent mouse. All experiments were performed in male *ad lib* fed C57Bl/6N mice during the light cycle.

Kapelczak, et al., Extended data Fig 5

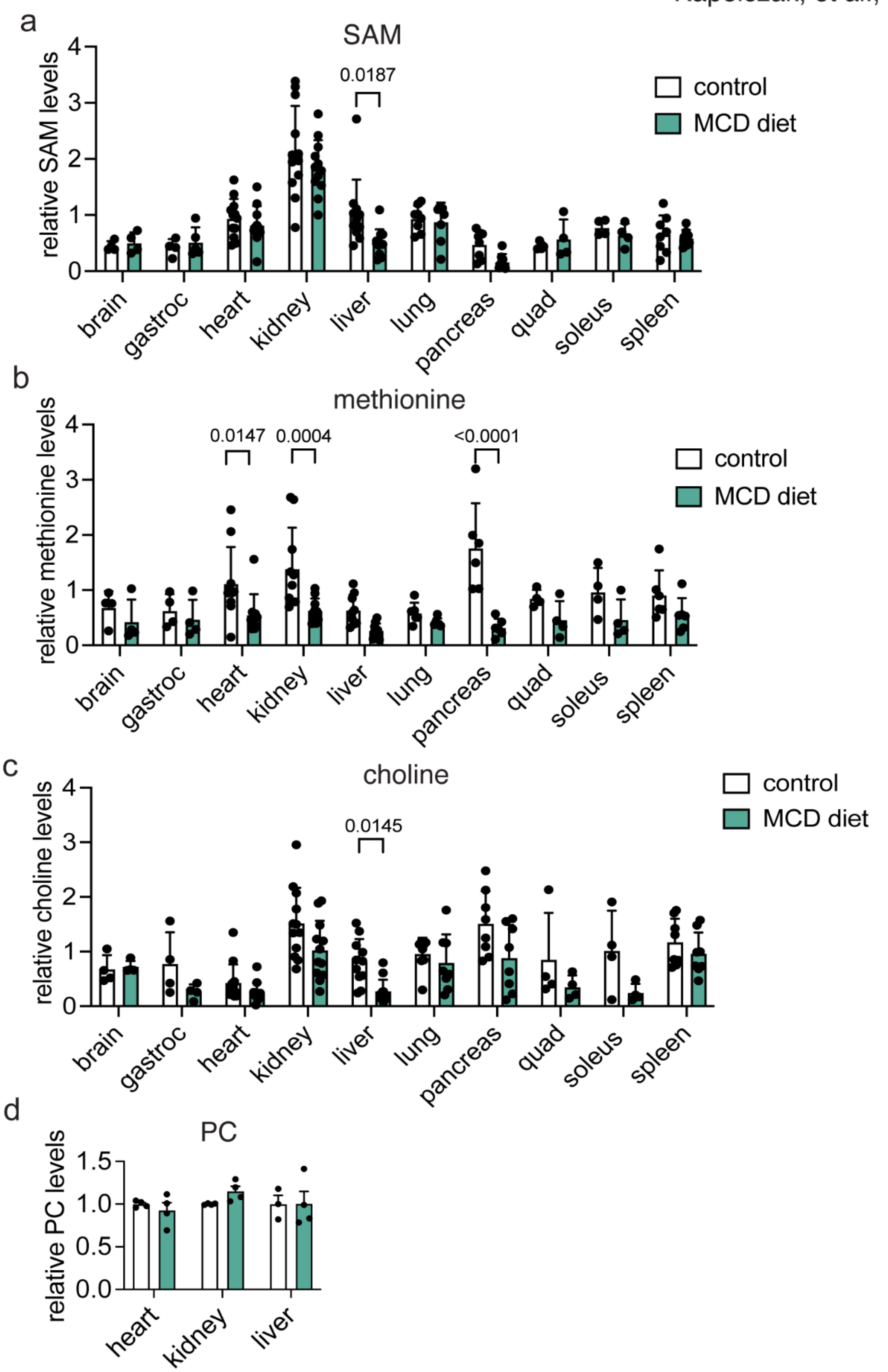


**Extended Data Figure 5. Changes in metabolite levels in tissue with dietary methionine and choline deficiency.**

(a) Relative ion counts for SAM across tissues in MCD diet and control diet fed mice.

(b) Relative ion counts for methionine across tissues in MCD diet and control diet fed mice.

(c) Relative ion counts for choline across tissues in MCD diet and control diet fed mice.

(d) Relative ion counts for pooled phosphatidylcholine (PC) species in tissues in MCD diet and control diet fed mice.

Bars represent mean $\pm$ S.D. P-value was calculated with mixed-effects analysis with Šídák correction for multiple comparisons. Each data point represents data from an independent mouse. All experiments were performed in male *ad lib* fed C57Bl/6N mice during the light cycle.

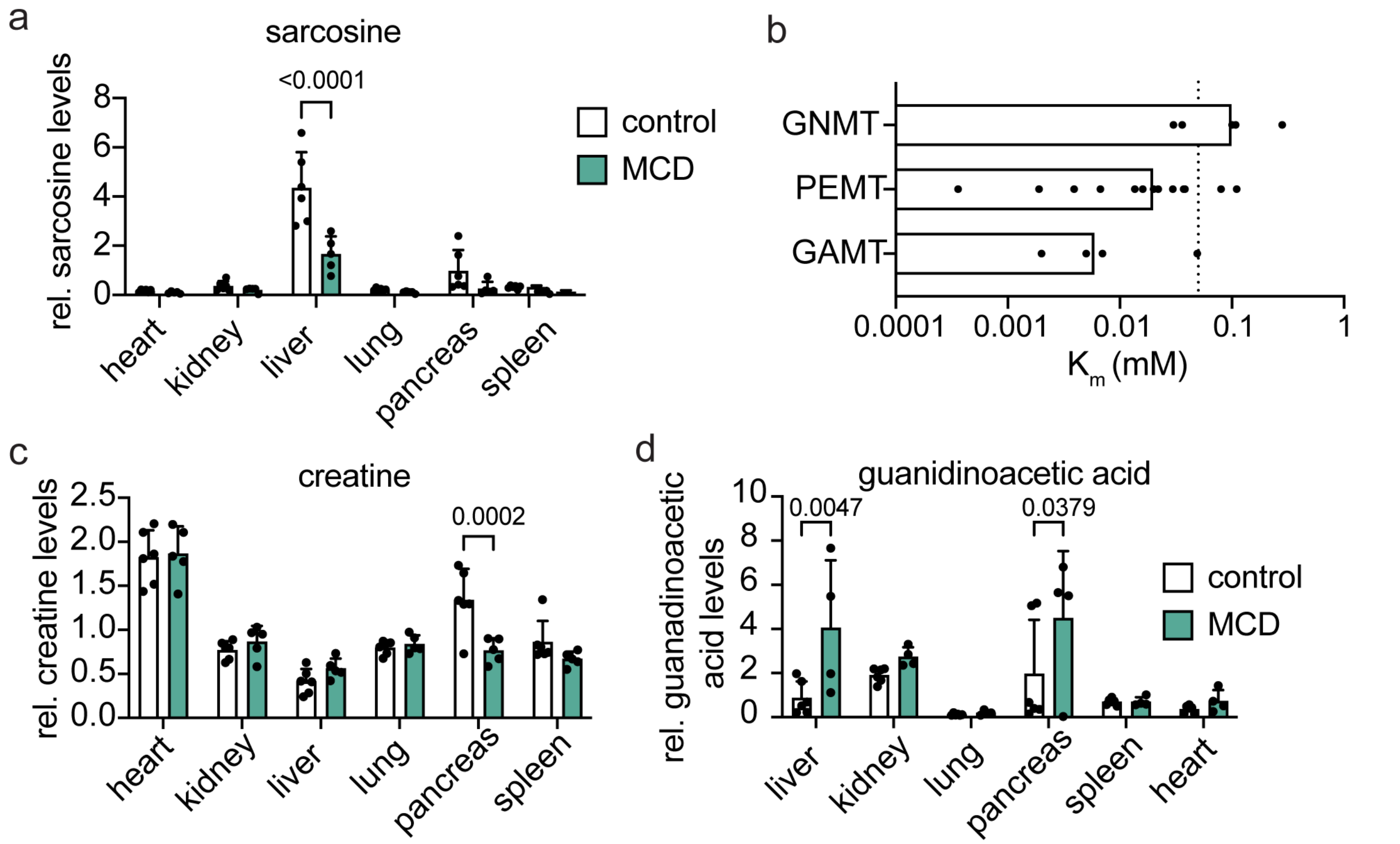


**Extended Data Figure 6. Moderate decreases in methylation in mice fed methionine and choline deficient diet.**

(a) Relative ion counts for sarcosine in tissues in MCD diet and control diet fed mice.

(b) Published $K_m$ values of mammalian methyltransferases for SAM as found in the Brenda database. Dotted line indicates the approximate physiological concentration of SAM in cells.

(c) Relative ion counts for sarcosine in tissues in MCD diet and control diet fed mice.

(d) Relative ion counts for guanidinoacetic acid in tissues in MCD diet and control diet fed mice

Bars represent mean ± S.D. P-value was calculated with mixed-effects analysis with Šídák correction for multiple comparisons. Each data point represents data from an independent mouse. All experiments were performed in male *ad lib* fed C57Bl/6N mice during the light cycle.

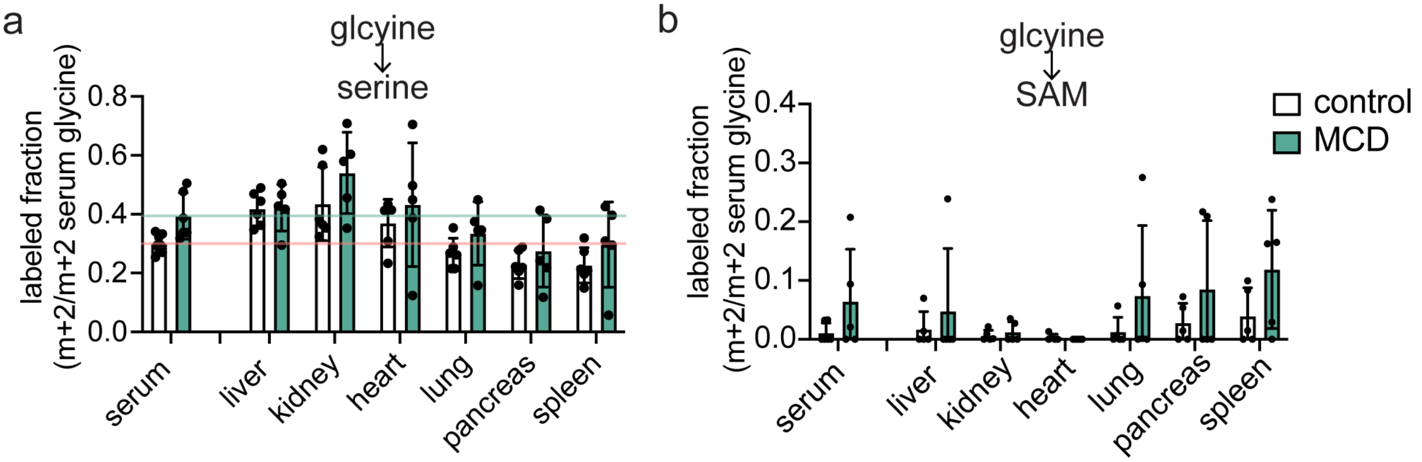


**Extended Data Figure 7. Glycine contribution to serine and SAM synthesis in control and MCD diet.**

(a) Labeling of tissue serine normalized to serum enrichment of glycine after 8 hour intravenous infusion of [U-$^{13}C_2$]glycine representing the fraction of serine that comes from glycine.

(b) Labeling of tissue SAM normalized to serum enrichment of glycine after 8 hour intravenous infusion of [U-$^{13}C_2$]glycine representing the fraction of serine that comes from glycine.

Bars represent mean ± S.D. Each data point represents data from an independent mouse. All experiments were performed in male *ad lib* fed C57Bl/6N mice during the light cycle.

Kapelczak, et al., Extended data Fig 8

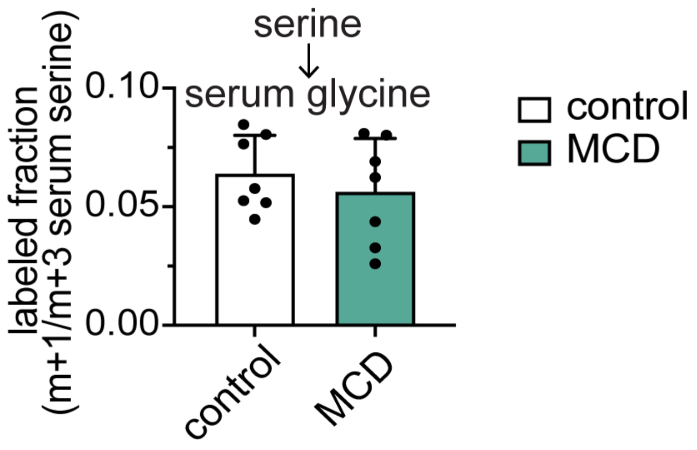


**Extended Data Figure 8. Fraction of glycine in serum that comes from serine.** Labeling of serum glycine normalized to serum enrichment of serine after 8 hour intravenous infusion of [2,3,3-$^{2}H_{3}$]serine.

Bars represent mean ± S.D. Each data point represents data from an independent mouse. All experiments were performed in male *ad lib* fed C57Bl/6N mice during the light cycle.
